# PV interneuron-targeted CRISPRa rescue of *SCN1A* haploinsufficiency in Dravet syndrome

**DOI:** 10.64898/2026.07.12.737793

**Authors:** Perry W.E. Spratt, Nicholas F. Trojanowski, Rajani M. George, Olivia Stevenson, Tessa Nottonson, Joshua Reiser, Lucia Capano, Andrew R. Field, Jaclyn Essig, Michael Faundo, Yan Hung, Navneet Matharu, Charlie Harper, Orrin Devinsky, Jordane Dimidschstein, Kathryn C. Allaway

## Abstract

Dravet syndrome is a severe epileptic encephalopathy caused by *SCN1A* haploinsufficiency, which leads to reduced Na_V_1.1 expression in parvalbumin (PV)-expressing interneurons and disrupted excitatory-inhibitory balance in the brain. We developed an AAV-based CRISPR activation system (AAV9-E2-dCas9-VP64) to selectively upregulate *SCN1A* from its endogenous locus in PV interneurons. An in vitro saturating guide RNA (gRNA) screen across the human *SCN1A* promoter identified a lead guide with robust and highly specific engagement of the *SCN1A* locus. This lead gRNA was validated in human Dravet syndrome model GABAergic neurons, where dose-dependent and specific *SCN1A* upregulation was observed. Intracerebroventricular (ICV) administration in a mouse model of Dravet syndrome produced dose-dependent improvement in survival as well as reduced susceptibility to hyperthermia-induced seizures and increased Na_V_1.1 protein expression, with maintained PV interneuron selectivity and minimal off-target expression. In a study in juvenile cynomolgus macaques, MRI-guided ICV administration of the vector was well tolerated, achieved broad cortical biodistribution, and maintained strong detargeting of peripheral tissues, with substantially lower peripheral dCas9 expression relative to the brain. These results support PV interneuron-selective *SCN1A* gene modulation via CRISPR activation as a promising therapeutic strategy for Dravet syndrome. AAV9-E2-dCas9-VP64 (RT101) is currently in preclinical development and is being advanced toward evaluation in the clinic.

---

Dravet syndrome (DS) is a severe, early-onset developmental and epileptic encephalopathy characterized by medication-resistant febrile and spontaneous seizures, developmental delay, and premature mortality due to sudden unexpected death in epilepsy (SUDEP).^1^ DS affects 1 in 15,700 individuals^2,3^ and over 80% of DS cases are caused by loss-of-function mutations in the gene *SCN1A* that result in haploinsufficiency of the voltage-gated sodium channel Na_V_1.1.^4^ Despite improvements in seizure management using anti-epileptic medications including stiripentol, cannabidiol, and fenfluramine,^5^ currently available treatments provide only partial control of seizures, lead to no or limited improvements in neurodevelopmental outcomes, and do not address the underlying genetic causes of the disorder.^6,7^ Thus, there remains a critical unmet need for disease-modifying therapies that meaningfully improve both seizure burden and neurodevelopmental outcomes.

Approaches that increase *Scn1a* expression in heterozygous *Scn1a* mouse models rescue seizure phenotypes and improve survival, even when expression is restored after symptom onset.^8–11^ Anti-sense oligonucleotides (ASOs) that target non-productive *SCN1A* splice variants to increase productive transcript expression have improved both seizure and neurodevelopmental endpoints in clinical trials,^12^ though this approach requires repeated intrathecal dosing under general anesthesia, a significant burden for the DS patient population. Alternatively, one-time dosing of an adeno-associated virus (AAV)-delivered zinc-finger transcriptional activator that increases the expression of the intact *SCN1A* allele has shown efficacy in both heterozygous *Scn1a* mouse models^13^ and DS patients.^14^ Together, these results validate endogenous *SCN1A*/Na_V_1.1 upregulation as a clinically viable therapeutic strategy for DS.

Na_V_1.1 is predominantly expressed in a subset of GABAergic interneurons and layer 5 excitatory neurons.^15–17^ Among these populations, parvalbumin (PV)-positive interneurons are most strongly implicated in DS seizure etiology. Na_V_1.1 is densely clustered in the axon initial segments of PV interneurons, the site of action potential initiation,^16^ and PV interneurons heterozygous for *Scn1a* exhibit impaired high-frequency action potential firing and propagation.^16,18–24^ Despite comprising only ~4% of cortical cells, selective knockout of *Scn1a* in PV interneurons is sufficient to reproduce both the seizure and cognitive impairment phenotypes observed in global heterozygous *Scn1a* models,^16,25–27^ demonstrating that dysfunction in this cellular population is a central driver of disease. Importantly, alteration of Na_V_1.1 expression in cellular populations that do not naturally express Na_V_1.1 has been shown to have negative effects on seizure and survival phenotypes.^17,28^

Despite the evidence that PV interneurons play a central role in DS, current disease-modifying therapies either lack cellular specificity entirely or rely on only broad targeting of GABAergic interneurons.^13,29^ There is therefore a possibility that the broad upregulation of Na_V_1.1 induced by these treatments could lead to alterations in network excitability, with potential implications for long-term safety and efficacy. In contrast, an intervention that upregulates Na_V_1.1 specifically in PV interneurons may enable restoration of inhibitory circuit function while minimizing perturbation of non-disease-relevant cells, and would thereby maximize therapeutic benefit while reducing potential risks associated with broader upregulation of Na_V_1.1 expression.

Here, we develop and evaluate an AAV9 vector that selectively upregulates *SCN1A* expression in PV interneurons. We use a CRISPR-activation (CRISPRa)-based approach, in which a catalytically deactivated Cas9 (dCas9) fused to a transcriptional activator is directed by a guide RNA (gRNA) to a target gene’s regulatory region, which boosts expression from the endogenous locus without editing the DNA sequence. Prior CRISPRa applications for DS and other haploinsufficient disorders that have been effective in mouse models^8,30,31^ have relied on either transgenic mouse lines or multi-AAV systems; here, we package all required components – a gRNA targeting the *SCN1A* promoter together with a Staphylococcus aureus dCas9-VP64 fusion driven by the PV interneuron-selective E2 enhancer element^32^ – within a single AAV9 vector. We show that this vector upregulates *SCN1A* in both mouse and human cells, restricts dCas9-VP64 expression to PV interneurons in vivo, and reduces seizure sensitivity and improves survival in a DS mouse model following a one-time intracerebroventricular (ICV) injection. ICV administration of the vector in non-human primates (NHPs) results in cortex-wide distribution of the vector and PV interneuron-specific transgene expression. These results demonstrate that PV interneuron-selective, CRISPRa-mediated *SCN1A* upregulation is a promising disease-modifying therapeutic approach for DS.

## Results

### Identification of a gRNA that upregulates murine *Scn1a* in vitro

Dravet syndrome poses unique opportunities and challenges for gene therapy approaches. 80% of DS cases are caused by loss-of-function variants in the *SCN1A* gene, and approaches that increase or restore *SCN1A* expression have been shown to be effective in animal models^8–11,13,28,29,33,34^ and in clinical trials.^12,14^ However, the large size of *SCN1A* (~6 kb coding sequence) exceeds the packaging capacity of adeno-associated virus (AAV) vectors,^35,36^ precluding conventional gene replacement strategies. Furthermore, the highly cell type-specific expression of Na_V_1.1 in PV-expressing GABAergic interneurons in the brain^16,25,37^ raises safety concerns of the overexpression of Na_V_1.1 in other cellular populations in the brain and throughout the body.^11,17^

To address these challenges, we devised a strategy for using CRISPRa to upregulate the endogenous *SCN1A* gene in heterozygotes from its native locus selectively in PV interneurons (Figure 1A), using a single AAV vector. This vector employs a catalytically deactivated *Staphylococcus aureus* Cas9 fused to the transcriptional activator VP64 (dCas9-VP64), which is guided to regulatory sequences of the *SCN1A* locus by a target-specific gRNA. This approach leverages the cell’s endogenous transcriptional machinery to increase *SCN1A* expression from the intact allele. PV interneuron selectivity is achieved using the E2 enhancer element to drive dCas9-VP64 expression. E2 is a cell type-specific regulatory element that restricts expression to PV interneurons,^32^ limiting the potential for off-target *SCN1A* modulation in other neuronal subtypes and minimizing transgene expression in peripheral tissues. While CRISPRa has been used extensively in mouse models to upregulate genes associated with haploinsufficient disorders, including *Scn1a*,^8,9^ in vivo phenotypic rescue using a CRISPRa system packaged entirely within a single AAV vector has not been demonstrated to date.

**Fig. 1.**
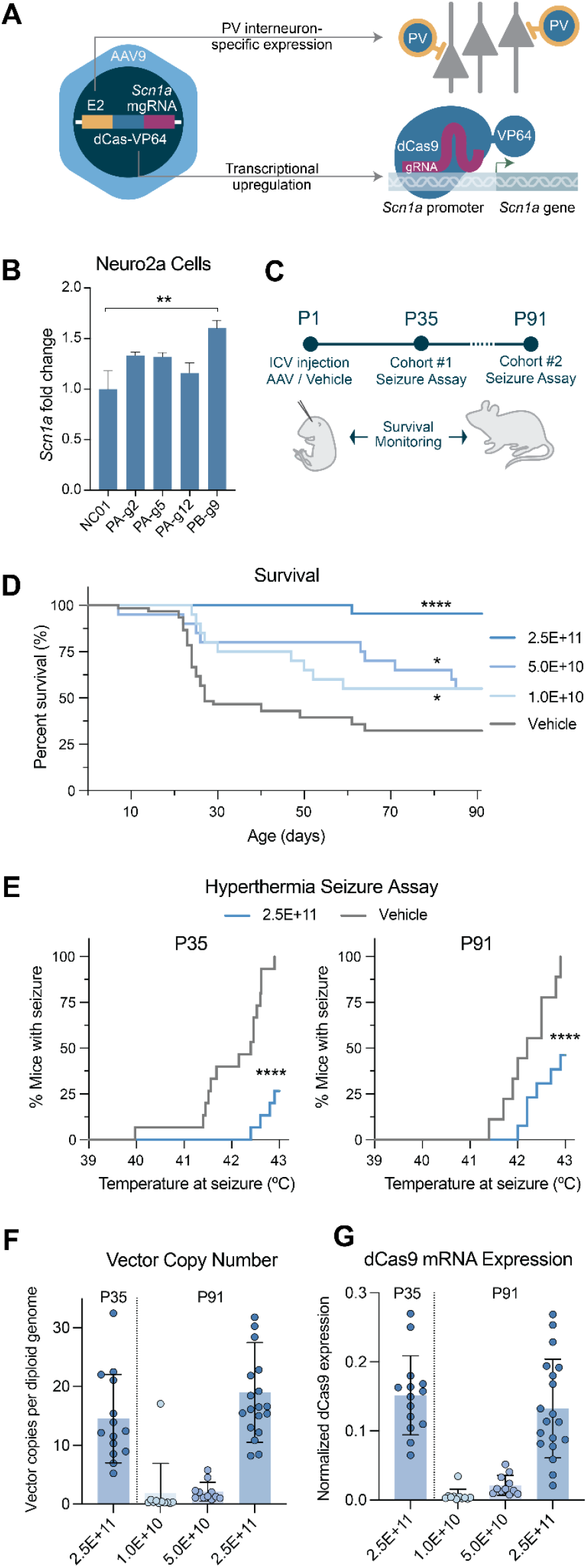
A PV interneuron-selective CRISPRa approach rescues survival and seizure susceptibility in *Scn1a*^+/-^ mice. **(A)** Schematic of the approach. The E2 regulatory element drives expression of dCas9-VP64 specifically in PV interneurons. Within the target cells, dCas9-VP64 associates with a sequence-specific gRNA to bind the *Scn1a* promoter, leading to upregulation of *Scn1a* expression. **(B)** Upregulation of *Scn1a* in vitro in Neuro2a cells with the top four mouse-specific gRNAs measured by qPCR following transfection with an E2-dCas9-VP64-hU6-gRNA all-in-one plasmid. Expression normalized to *Actb* and shown as fold change relative to non-targeting gRNA (NC01). n=2 per gRNA. Statistical significance determined by one-way ANOVA with Dunnett’s multiple comparisons relative to NC01, **p=0.0059. See also Figure S1. **(C)** Design of in vivo study. Mice were treated with either vehicle or our vector containing AAV9-E2-dCas9-VP64-*Scn1a*-PB-g9 gRNA at P1, then a subset of mice were subjected to hyperthermia at either P35 or P91. **(D)** Percent survival of *Scn1a*^+/-^ mice following P1 ICV injections (Vehicle: n=60; 2.5E+11 vg: n=45, p<0.0001; 5.0E+10 vg: n=20, p=0.031; 1.0E+10 vg: n=20, p=0.034, log-rank test). **(E)** Percentage of mice that seized at or below the indicated temperature. Left: mice assayed at P35, n=15 mice per group, ****p<0.0001, log-rank test. Right: mice assayed at P91, Vehicle, n=9; 2.5E+11, n=12, ****p<0.0001, log-rank test. **(F)** Vector copies per diploid genome from mouse somatosensory cortex at P35 and P91. A custom U6-targeting assay was used to quantify vector genomes, and a Ptbp2-targeting assay was used to quantify mouse genomes. Each point represents one mouse, bars represent group means, error bars represent standard deviation. P35: n=14; 1.0E+10: n=11; 5.0E+10: n=11; 2.5E+11: n=18. **(G)** dCas9 mRNA expression normalized to *Hprt* mRNA expression at P35 and P91. Each point represents one mouse, bars represent group means, error bars represent standard deviation. P35: n=14; 1.0E+10: n=11; 5.0E+10: n=11; 2.5E+11: n=19. See also Figure S2.

We validated this approach in a proof-of-concept study using a mouse model of DS. The murine *Scn1a* locus has well defined gene regulatory regions including distal and proximal promoters and an enhancer region within the first intron^32,38,39^. gRNAs targeting these regulatory sequences were screened in vitro in Neuro2a cells by inserting them into a plasmid expressing dCas9-VP64 ubiquitously under the CMV promoter (Figure S1A). Top performing guides were subsequently validated with a plasmid in which dCas9-VP64 was controlled by the E2 regulatory element (Figure 1B). A gRNA targeting the proximal promoter (*Scn1a*-PB-g9) was selected for in vivo validation on the basis of robust *Scn1a* upregulation and an absence of modulation of the expression of neighboring genes (Figure S1B). A single plasmid containing both the E2-dCas9-VP64 cassette and the *Scn1a*-PB-g9 gRNA was then packaged into an AAV9 vector for in vivo efficacy testing.

### PV interneuron-targeted CRISPRa upregulation of *Scn1a* rescues core phenotypes in a Dravet mouse model

The efficacy of this vector was next tested in a Dravet mouse model (*Scn1a*^*tm1Kea*^), which exhibits premature mortality and is susceptible to hyperthermia-induced seizures, recapitulating both the sudden unexpected death in epilepsy (SUDEP) and febrile seizure features of DS.^40,41^ Our vector was administered to postnatal day (P)1 *Scn1a*^+/-^ mice via bilateral ICV injection at three dose levels: 1.0E+10, 5.0E+10, and 2.5E+11 vector genomes (vg) per mouse. A cohort of vehicle-treated *Scn1a*^+/-^ mice served as controls. Animals were monitored daily for survival through P91, and a subset of control and high-dose animals were assessed for hyperthermia-induced seizure sensitivity at P35 and P91 (Figure 1C).

Treatment with our vector produced dose-dependent improvements in survival, with all doses showing significant improvement relative to vehicle-treated controls, and only one mortality out of 40 mice treated at the highest dose (Figure 1D). Body weights did not differ significantly between treatment groups over the course of the study.

Seizure threshold was evaluated in vehicle-treated and high-dose (2.5E+11 vg) *Scn1a*^+/-^ mice by subjecting subsets of animals to a hyperthermia-induced seizure protocol at P35 and P91. At P35 all assessed vehicle-treated animals exhibited seizures, whereas only 27% of mice treated with the highest dose seized during the assay (Figure 1E, left). The protective effect was maintained at P91, when all vehicle-treated animals again exhibited seizures compared to only 46% of high-dose animals (Figure 1E, right). Additionally, the average temperature at seizure was higher in treated animals compared to untreated animals. Overall, these results demonstrate durable seizure protection through at least three months following dosing.

Comprehensive molecular and histological analyses were performed on brain tissue collected at P35 (in animals subjected to the hyperthermia protocol) and at P91 (all surviving animals). As expected, vector copy number (vector genome copies per diploid genome, VCN) and dCas9-VP64 mRNA expression increased with dose in cortical samples assessed at P91 (Figure 1F, G). Vector persistence and stability of transgene expression were assessed in high-dose animals at P35 and P91, and no significant differences in either VCN or dCas9 expression between ages were observed.

At P35 and P91, *Scn1a* mRNA expression trended toward upregulation in treated animals, although differences did not reach statistical significance (Figure S2A, B). In contrast to the restricted expression pattern of Na_V_1.1, *Scn1a* mRNA is expressed broadly,^15^ therefore PV interneuron-specific *Scn1a* upregulation is unlikely to be detectable in measurements of bulk *Scn1a* transcript levels. The observed effect on NaV1.1 protein expression was larger, likely due to the more restricted expression of Na_V_1.1 protein to PV interneurons compared to other neuronal cell types, though statistical significance was only observed at P35 (Figure S2C, D).

### Saturating screen of the human *SCN1A* locus identifies a highly efficacious and specific gRNA

We next sought to develop and validate a version of the vector suitable for clinical development, as our PB-g9 gRNA does not have a binding site in the human *SCN1A* promoter region. To this end, we nominated and tested gRNAs targeting the human *SCN1A* proximal promoter, since targeting this promoter led to upregulation and phenotypic improvement in the mouse model. In silico nomination was performed to saturate the locus from 1000 bp upstream to 500 bp downstream of the transcription start site (TSS), leading to the nomination of 26 candidate gRNAs (Figure 2A).

**Fig. 2.**
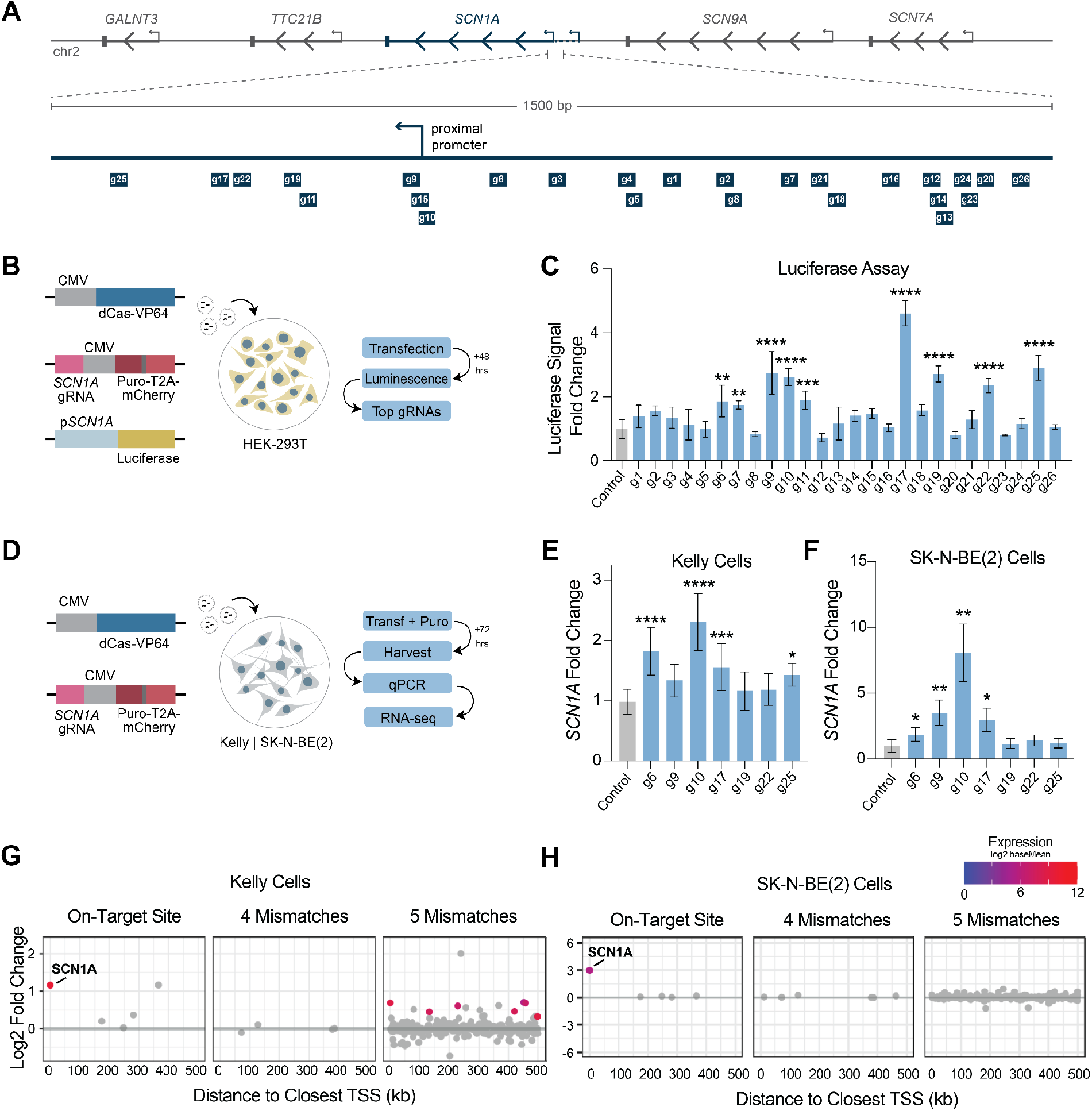
A saturating gRNA screen to identify a potent and highly specific gRNA targeting *SCN1A* in the human genome. **(A)** Location of gRNAs tested. **(B)** Luciferase assay: HEK-293T cells were transfected with the three plasmids indicated. **(C)** Normalized luminescence (fold change relative to control). n=3 per gRNA, results from three non-targeting gRNAs were combined for control. Significance determined by one-way ANOVA with Dunnett’s multiple comparisons. g6: p=0.0015; g7: p=0.0099; g11: p<0.001; g9, g10, g17, g19, g22, g25: p<0.0001. **(D)** Dual vector system used for screening of individual *SCN1A*-targeting gRNAs in cell lines (Kelly & SK-N-BE(2)). **(E, F)** *SCN1A* mRNA expression fold change relative to non-targeting gRNA control in Kelly **(E)** or SK-N-BE(2) **(F)** cells determined by probe-based qPCR and normalized to *RPS18* (Kelly) or the geometric mean of *RPS18, HPRT*, and *18SN5* (SK-N-BE(2)) expression. n=6 per gRNA. Significance determined by one-way ANOVA with Dunnett’s multiple comparisons. Kelly – g25: p=0.0103, g17: p<0.001, g6, g10: p<0.0001. SK-N-BE(2) – g6: p=0.0343; g9: p=0.0052, g10: p=0.0029, g17: p=0.0109. **(G**,**H)** Off-target analysis in Kelly **(G)** and SK-N-BE(2) **(H)** cells. Genes with normalized counts ≥10 in ≥2 samples and a TSS within 500 kb of a predicted *SCN1A*-g10 on- or off-target binding site (≤5 mismatches) were included. The x-axis indicates the distance from each predicted binding site to each gene’s nearest annotated TSS. *SCN1A*-g10 has no predicted off-target sites with 1-3 mismatches. Significantly upregulated (log2FC>0.3, pAdj<0.05) genes are shown in color (indicating baseMean expression); gray dots do not reach significance/expression thresholds.

First, each gRNA was tested using a luciferase assay in HEK-293T cells in which the *SCN1A* promoter was cloned upstream of firefly luciferase and co-transfected with the dCas9 and gRNA-containing plasmids (Figure 2B). Luminescence was then measured to assess the degree of upregulation of the luciferase gene triggered by gRNA binding activity on the *SCN1A* promoter. Several gRNAs elicited significant luciferase activity, and the top-performing gRNAs were selected for validation and further testing (Figure 2C).

Each gRNA was then cloned into a dual vector screening system in which CMV-dCas9-VP64 was expressed from one plasmid, while each gRNA was expressed from a second plasmid that also included a puromycin selection cassette to allow for enrichment of transfected cells. These top gRNAs were then tested in Kelly and SK-N-BE(2) neuroblastoma cell lines for upregulation of endogenous *SCN1A*, using qPCR to assess *SCN1A* mRNA expression relative to a non-targeting gRNA control (Figure 2D). Of these, *SCN1A*-g10 showed the most potent *SCN1A* upregulation in both cell lines (Figure 2E, F).

To ensure suitability for use in a clinical product, the genomic off-target profile of *SCN1A*-g10 was thoroughly interrogated. First, CasOFFinder^42^ was used to perform *in silico* off-target analysis to identify all potential off-target binding sites in the human genome with up to five mismatches or one DNA or RNA bulge. *SCN1A*-g10 had no predicted off-target binding sites with up to three mismatches, three predicted binding sites with four mismatches, and 70 sites with five mismatches. To assess whether any of the putative off-target sites were associated with transcriptional activation of nearby genes, we performed bulk RNA-seq on samples from Kelly and SK-N-BE(2) cells and evaluated the expression of genes with a transcriptional start site (TSS) within 500 kb of each putative binding site (Figure 2G,H).

At the *SCN1A* locus, expression of all neighboring genes did not differ significantly between cells treated with *SCN1A*-g10 and those treated with the non-targeting control gRNA, with the exception of *SCN1A* itself. Likewise, among predicted off-target sites with up to four mismatches, no genes meeting the distance and expression thresholds were significantly upregulated in either cell line. At five mismatches, seven genes were significantly upregulated in Kelly cells. However, none of these were significantly upregulated in SK-N-BE(2) cells. Given the low likelihood of bona fide dCas9 binding at sites with five or more mismatches, this suggests that the *SCN1A*-g10 gRNA has an exceptionally favorable off-target profile.

### CRISPRa upregulation of *SCN1A* in human DS model neurons

To verify the activity of the *SCN1A*-g10 gRNA and CRISPRa approach in a human disease model, we tested these components in DS model (heterozygous loss-of-function mutation in *SCN1A*) iPSC-derived GABAergic neurons in vitro. A surrogate system was used to deliver the CRISPRa components efficiently in this in vitro system that differs from our in vivo product in two ways: 1) the PHP.eB capsid was used due to inefficient AAV9 transduction of neurons in vitro and 2) a PGK promoter was used to drive expression of dCas9 due to the relatively immature state of GABAergic neurons in vitro (Figure 3A).

**Fig. 3.**
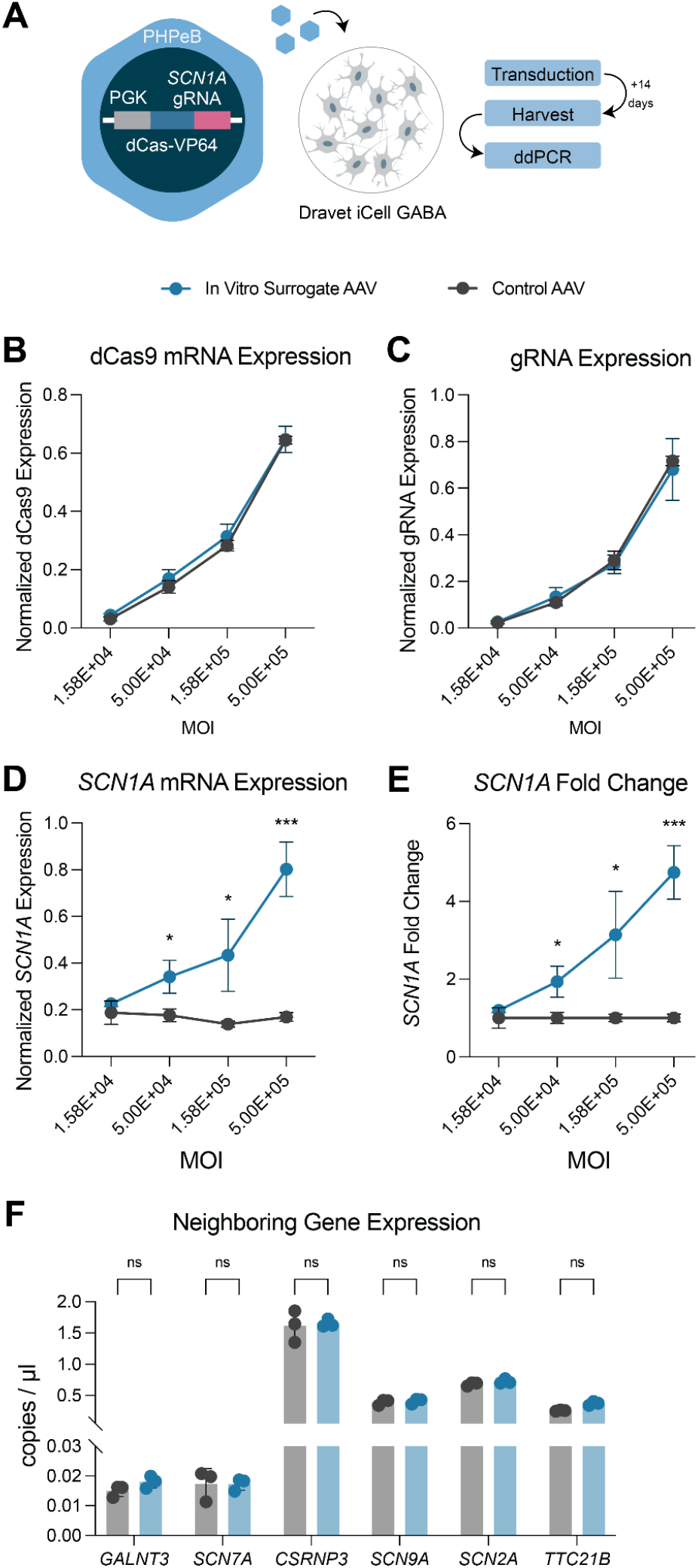
CRISPRa upregulation of *SCN1A* in human DS model neurons. **(A)** The PGK promoter was used to drive dCas9-VP64 expression in the same vector as the hU6-*SCN1A*-g10 gRNA cassette and packaged in AAV-PHP.eB for efficient in vitro expression and used to transduce iCell GABA Dravet neurons. For control virus, the *SCN1A*-gRNA was replaced with a non-targeting gRNA. n=3 per MOI. **(B)** dCas9 mRNA expression in both control AAV-treated and *SCN1A*-targeting AAV-treated neurons at each MOI. **(C)** gRNA expression at each MOI. **(D)** *SCN1A* mRNA expression normalized to the geometric mean of *RPS18* and *TBP expression*. **(E)** *SCN1A* mRNA expression fold change relative to the average of control samples at each MOI. For **(D)** and **(E)**, statistical significance was determined by multiple unpaired t-tests. 5.00E+04: p=0.019749. 1.58E+05: p=0.030001. 5.00E+05: p=0.000744. **(F)** mRNA expression of genes neighboring *SCN1A* in the 5.00E+05 MOI condition for control (gray) or treated (blue) samples. Multiple unpaired t-tests were performed with no statistically significant comparisons. Expression was measured by ddPCR. Error bars indicate standard deviation. MOI: multiplicity of infection.

Four multiplicities of infection (MOIs) were used to assess dose-responsive upregulation compared to a control AAV containing non-targeting gRNA by ddPCR following 14 days in culture post-transduction. dCas9 expression and gRNA expression were highly consistent at all MOIs across the non-targeting control and the *SCN1A*-targeting AAV (Figure 3B, C). *SCN1A* expression was upregulated in a dose-dependent manner relative to the non-targeting control following treatment with the *SCN1A*-targeting AAV (Figure 3D, E). None of the neighboring genes within the vicinity of *SCN1A* (*GALNT3, SCN7A, CSRNP3, SCN9A, SCN2A*, and *TTC21B*) were differentially expressed (Figure 3F), suggesting that activity of the gRNA also remained specific to *SCN1A* in this cellular context.

### Treatment with AAV9-E2-dCas9-VP64-*SCN1A*-g10 restores Na_**V**_1.1 levels and rescues premature lethality in mice in a dose-dependent manner

Although it was not included in our initial mouse gRNA screen, *SCN1A*-g10 has a conserved binding site in the mouse genome. To test if this gRNA can also drive mouse *Scn1a* upregulation, we tested it in our luciferase and in vitro screening pipeline as described above. To our surprise, we found that *SCN1A*-g10 outperformed *Scn1a*-PB-g9 and all other previously screened guides in our luciferase screen (Figure S3A) and in Neuro2a cells (Figure S3B). Next, we tested the activity of our *SCN1A*-g10 gRNA in vivo in *Scn1a*^+/-^ mice. As in our proof-of-concept experiment with the mouse gRNA *Scn1a*-PB-g9, our vector with *SCN1A*-g10 was administered to P1 mice via bilateral ICV injection at three dose levels: 5.0E+10, 1.0E+11, and 2.5E+11 vg per mouse (Figure 4A). We again observed a dose-dependent increase in survival at P35, with 100% survival achieved with a dose of 2.5E+11 vg (Figure 4B). Underlying this survival was a dose-dependent upregulation of *Scn1a* (27% at 2.5E+11 vg, Figure 4C) and Na_V_1.1 (35% at 2.5E+11 vg, Figure 4D). There were no differences in body weight across any doses over the course of the study, indicating that AAV9-E2-dCas9-VP64-*SCN1A*-g10 is generally well tolerated.

**Fig. 4.**
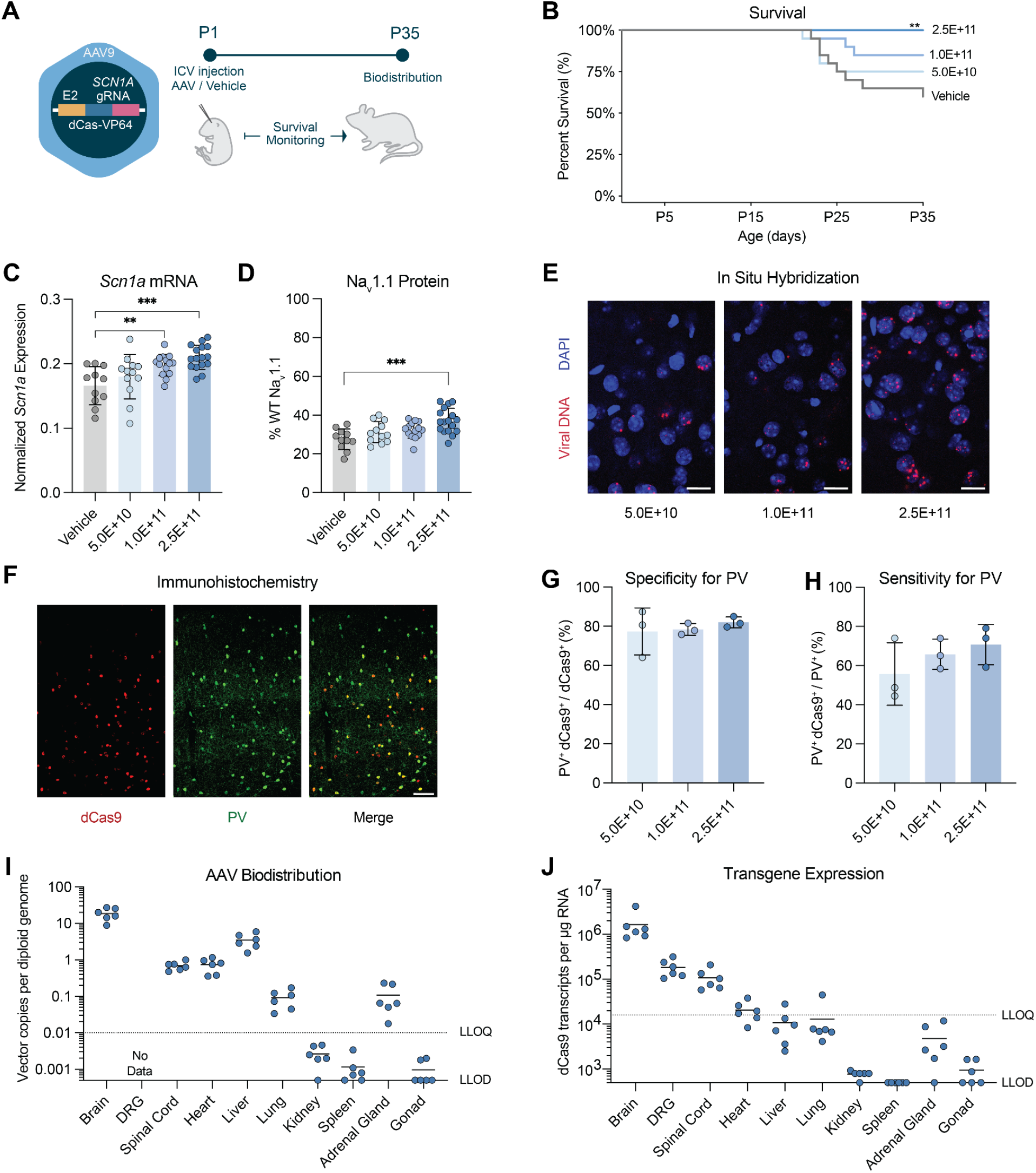
Dose-dependent AAV9-E2-dCas9-VP64 treatment restores Na_V_1.1 levels and rescues premature lethality in mice. **(A)** Mice were treated with either vehicle or our vector containing AAV9-E2-dCas9-VP64-*SCN1A*-g10 gRNA at P1 and sacrificed at P35. See also Figure S3. **(B)** Percent survival of *Scn1a*^+/-^ mice following P1 ICV injections. **p=0.0017, individual logrank tests vs. vehicle; n=20 per group. **(C)** *Scn1a* mRNA is upregulated by AAV9-E2-dCas9-VP64-*SCN1A*-g10 in *Scn1a*^+/-^ mice. *Scn1a* mRNA normalized to *Hprt* mRNA. Each point represents one mouse somatosensory cortex. Each bar represents the mean of all mice treated with the indicated dose, error bars represent standard deviation. **p=0.0049, ***p=0.0001, Holm-Šídák’s multiple comparisons test following one-way ANOVA. Vehicle: n=11; 5.0E+10: n=12; 1.0E+11: n=14; 2.5E+11: n=16. **(D)** Na_V_1.1 is upregulated by AAV9-E2-dCas9-VP64-*SCN1A*-g10 in *Scn1a*^+/-^ mice. Na_V_1.1 levels are shown as a percentage of average Na_V_1.1 measured in littermate WT controls. Each point represents one mouse somatosensory cortex. Each bar represents the mean of all mice treated with the indicated dose, error bars represent standard deviation. *** p=0.0001, Holm-Šídák’s multiple comparisons test following one-way ANOVA. Vehicle: n=11; 5.0E+10: n=12; 1.0E+11: n=14; 2.5E+11: n=17. **(E)** Representative fluorescence microscopy images showing dose-dependent increases in viral DNA puncta detected by RNAscope (red) with nuclei stained by DAPI (blue). Scale bar (20 µm) and imaging conditions were kept constant across all panels. **(F)** Representative fluorescence microscopy images showing immunohistochemical validation of dCas9-VP64 (red, left panel) expression in PV-positive neurons (green, middle panel), with co-localization demonstrated in the merged overlay (right panel). Scale bar represents 100 µm. **(G)** Specificity of dCas9-VP64 expression for PV-positive neurons across doses in somatosensory cortex, quantified as the percentage of dCas9-positive cells that are co-positive for PV. Data are presented as mean ± SD; individual data points represent the average of three sections per assessed mouse (n=3 per group). **(H)** Sensitivity of dCas9-VP64 expression among PV-positive neurons across doses in somatosensory cortex, quantified as the percentage of PV-positive cells that are co-positive for dCas9. Data are presented as mean ± SD; individual data points represent the average of three sections per assessed mouse (n=3 per group). **(I)** Vector copies per diploid genome across brain and peripheral tissues. Each point represents the measurement from one mouse from the 2.5E+11 dose, n=6. Dashed lines denote LLOQ and LLOD. **(J)** dCas9 transcripts per µg RNA across brain and peripheral tissues. Each point represents the measurement from one mouse from the 2.5E+11 dose, n=6. Dashed lines denote LLOQ and LLOD.

As expected, in situ hybridization assessment of vector genome DNA revealed dose-dependent increases in vector load in somatosensory cortex, with an increasing number of DNA puncta observed per nucleus with increasing dose (Figure 4E). To confirm that E2-driven expression maintained cell type selectivity across doses, we performed immunohistochemistry assessment of dCas9-VP64 within the same cortical region (Figure 4F). We observed a high specificity for PV interneurons across doses, with ~80% of dCas9-VP64 positive cells co-expressing PV (Figure 4G). The proportion of PV interneurons expressing detectable dCas9-VP64 increased with dose, rising from 45.4±23.7% at the lowest dose to 73.9±9.1% at the highest dose (Figure 4H).

To understand the peripheral biodistribution and activity of AAV9-E2-dCas9-VP64-*SCN1A*-g10 following ICV injection, we quantified vector genome abundance and dCas9 mRNA expression across numerous peripheral organs as well as the spinal cord. Vector genomes were detectable in the liver, heart, spinal cord, lung, and adrenal gland (Figure 4I), as expected from the known tropism of AAV9.^43–45^ Encouragingly, we found that dCas9 mRNA was only consistently detectable in the spinal cord and DRG (Figure 4J). Given the high vector biodistribution detected in the liver, the >100-fold lower dCas9 mRNA expression in the liver than the cortex demonstrates that AAV9-E2-dCas9-VP64-*SCN1A*-g10 effectively detargets dCas9 expression from the liver. Further, the expression of dCas9 mRNA in the spinal cord and DRG was >9x lower than that in the brain, underscoring the fact that even within the nervous system, E2 confers a high degree of specificity.

### AAV9-E2-dCas9-VP64-*SCN1A*-g10 is well tolerated and detargets peripheral organs in non-human primates

In order to proceed with AAV9-E2-dCas9-VP64-*SCN1A*-g10 as a potential clinical product, we sought to understand its safety and biodistribution profile following ICV injection in NHPs. To examine this, we performed a 42-day study in six juvenile cynomolgus macaques (two male, four female) with unilateral MRI-guided administration using the clinically relevant Clearpoint cannula system (Figure 5A, B). Two doses (1.9E+13 and 6.0E+13 vg per animal) were selected based on efficacy observed in mouse studies and allometric scaling of brain weights across species. No steroids or immunosuppressants were administered.

**Fig. 5.**
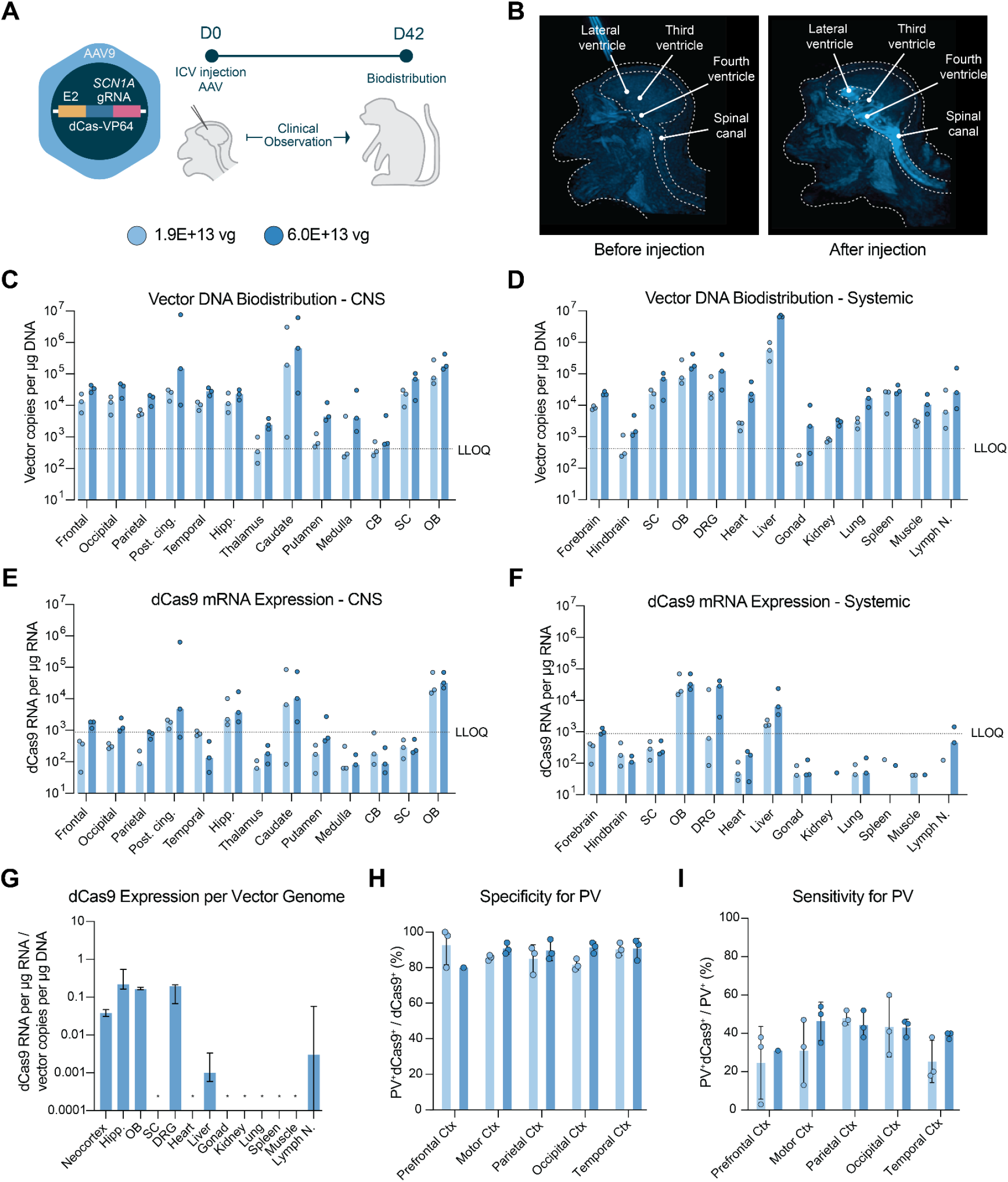
AAV9-E2-dCas9-VP64-*SCN1A*-g10 is well tolerated and detargets peripheral organs in NHP. **(A)** Juvenile cynomolgus macaques were treated with our vector containing AAV9-E2-dCas9-VP64-*SCN1A*-g10 and observed for 42 days before sacrifice. **(B)** MRIs showing the flow of contrast agent through the ventricular system during ICV injection. **(C)** Vector biodistribution across CNS regions. Each bar represents the median value for three animals. Each point represents the median value across hemispheres and punches for one animal. The dashed line represents the LLOQ. **(D)** Systemic vector biodistribution. Each bar represents the median value for three animals. Each point represents the median value for one animal. The dashed line represents the LLOQ. **(E)** dCas9 mRNA expression across CNS regions. Each bar represents the median value for three animals. Each point represents the median value across hemispheres and punches for one animal. The dashed line represents the LLOQ. **(F)** Systemic dCas9 mRNA expression. Each bar represents the median value for three animals. Each point represents the median value for one animal. The dashed line represents the LLOQ. **(G)** dCas9 mRNA expression per vector genome. This ratio is calculated by dividing the dCas9 mRNA expression per µg RNA input by the vector genome copies per µg DNA input. Each bar represents the median value across three animals; error bars denote the range; asterisks indicate values below the LLOQ. Data in panels C-G were quantified by ddPCR. **(H, I)** Immunohistochemical analysis of specificity **(H)** and sensitivity **(I)** of dCas9-VP64 expression for PV-positive neurons across doses in different cortical regions, quantified as the percentage of dCas9-positive cells co-positive for PV (specificity) or PV-positive cells co-positive for dCas9 (sensitivity). Data are presented as mean ± SD; individual data points represent the combined counts from two ROIs per region (n=3 per group). For the prefrontal cortex at the high dose (6.0E+13 vg), only one animal yielded sections suitable for quantification (n=1).

Administration of AAV9-E2-dCas9-VP64-*SCN1A*-g10 was generally well tolerated. There were no treatment-related changes in body weight and no significant adverse effects noted throughout the duration of the study. In one animal, intraoperative MRI scans showed a contrast agent spreading along the longitudinal fissure, suggesting that the test article in that animal leaked out of the ventricle along the injection track. This animal showed localized inflammation in the cingulate cortex adjacent to the longitudinal fissure. All other histopathological findings were otherwise considered minimal to moderate and were consistent with expected findings following AAV administration.

To measure vector biodistribution and activity, DNA and RNA were extracted from various tissues. With the exception of the animal in which the test article leaked, ddPCR analysis of brain samples revealed consistent vector biodistribution across cortical regions, with generally lower biodistribution in regions of the midbrain and hindbrain (Figure 5C). Vector biodistribution was detected in many peripheral tissues, and followed a similar pattern of AAV9 biodistribution observed by other groups^43–45^ with a much higher amount of vector present in the liver than in other organs (Figure 5D). The vector was also detected in other regions of the central nervous system (spinal cord, olfactory bulb) and peripheral nervous system (DRG), as expected from ICV injections of AAV9. This demonstrates that we are able to achieve uniform brainwide AAV biodistribution using a ClearPoint MRI-guided AAV administration system similar to one that can be used for dosing patients, without unexpected off-target biodistribution.

E2 has previously been reported to target PV interneurons with ~90% specificity in the macaque,^32^ but its activity in other tissues has not been assessed in detail. We found that in the high dose (6.0E+13 vg) group, dCas9 mRNA expression in the brain followed the same trends as the vector biodistribution, with generally homogeneous expression across cortical regions and less expression in most subcortical regions (Figure 5E). dCas9 expression was also detected in other regions of the nervous system, including the olfactory bulb and DRG. In the low dose (1.9E+13 vg) group, dCas9 expression was generally below our limit of quantitation (Figure 5F). The only tissue outside of the nervous system where dCas9 was expressed at quantifiable levels was the liver, where the vector-normalized dCas9 mRNA expression was >35x lower than in the brain, confirming the high degree of specificity of E2 across species.

Further supporting the specificity of vector expression for PV interneurons, immuno-histochemical analysis across five cortical regions (prefrontal, motor, parietal, occipital, and temporal cortex) showed that dCas9 protein expression was 80-93% specific for PV cells at both doses (Figure 5H; low dose: 82-93%, high dose 80-91%). Notably, the fraction of PV interneurons expressing dCas9 (i.e., sensitivity) varied across regions but was similar between doses, ranging from 25-48% at the low dose and 31-46% at the high dose (Figure 5I). These findings suggest that transgene expression saturation may already be achieved at the lower dose in several cortical regions, although modest increases in sensitivity were observed at the higher dose in motor and temporal cortex. Together, these data demonstrate that E2 can effectively drive dCas9 expression in PV interneurons throughout the primate brain while substantially detargeting tissues outside the nervous system.

## Discussion

In this study, we demonstrated that an AAV9-delivered, E2-driven CRISPR activation system enables highly selective upregulation of *SCN1A* in PV interneurons, leading to the rescue of key clinically relevant phenotypes in a mouse model of Dravet syndrome. Following initial proof-of-concept using a mouse surrogate vector, a gRNA targeting human *SCN1A* was identified through a saturating screen. This optimal gRNA drove robust and highly specific activation of *SCN1A* in multiple neuronal models, including human DS model GABAergic neurons. The final vector was then validated in a DS mouse model, in which we performed extensive characterization of biodistribution and transgene expression. Biodistribution and cell type specificity were further evaluated in NHPs, demonstrating maintenance of PV interneuron-restricted expression across species. Collectively, these findings support the feasibility of a PV interneuron-restricted, CRISPRa-based approach for targeted *SCN1A* upregulation in DS and establish a foundation for further translational development.

A key question underlying this work was whether selective restoration of *SCN1A* expression in PV interneurons is sufficient to drive meaningful phenotypic rescue in DS. Despite representing only ~4% of cortical cells, PV interneurons play a central role in maintaining excitatory-inhibitory balance, and their dysfunction is directly related to DS. While *SCN1A* mRNA is expressed broadly within the brain, Na_V_1.1 protein is enriched in the axon initial segments of cortical and hippocampal PV interneurons, where it plays a key role in action potential initiation and propagation.^16,19–24^ Consistent with this, selective deletion of *Scn1a* in PV interneurons recapitulates core phenotypes observed in global haploinsufficient models.^27^

However, other populations of neurons, including somatostatin (SST)- and vasoactive intestinal peptide (VIP)-expressing interneurons and layer 5 pyramidal neurons, also have reported roles in DS pathology,^15,19,46,47^ leaving open the question of whether restoration only in PV interneurons is sufficient to rescue DS phenotypes. The results here suggest that the phenotypic improvements achieved with a PV interneuron-selective approach are comparable to those using less specific rescue strategies^8,13,28,29^ and support the use of more stringent, cell type-specific regulatory elements to maximize therapeutic benefit while minimizing potential off-target effects. Notably, the E2 promoter used in this study is derived from the first intron of *SCN1A* and is likely a native *SCN1A* enhancer. The small fraction of non-PV cells that express the dCas9 transgene may therefore represent some of these other small neuronal populations relevant to DS,^32^ raising the possibility that this promoter is uniquely well suited to the underlying biology of the disease.

A defining feature of the approach reported here is the incorporation of multiple layers of specificity within the vector design, supporting its potential suitability for clinical development. First, delivery via AAV9 through ICV administration results in preferential transduction of neuronal populations relative to other serotype and route combinations. Although AAV9 can distribute to peripheral tissues following ICV delivery, with greater peripheral exposure observed in primates compared to mice, this approach represents a clinically validated strategy for achieving broad CNS transduction with relative detargeting of peripheral organs.^13,45,48–51^ Second, the E2 promoter restricts expression of the transgene to a key cell type implicated in DS and is sufficient to lead to therapeutic benefit in a mouse model of the disease. Importantly, this specificity for PV interneurons is maintained in NHPs and, together with the homogeneous transgene expression observed across brain regions, supports the translational potential of this vector. Finally, the CRISPRa-mediated approach to *SCN1A* upregulation exhibited high genomic specificity, supported by minimal predicted off-target binding and no detectable transcriptional activation of genes near predicted off-target sites.

Taken together, these features distinguish this approach from other disease-modifying strategies in development for DS. Antisense oligonucleotides (ASOs) are the most clinically advanced disease-modifying approach for DS and have demonstrated that upregulation of *SCN1A* in DS patients leads to significant therapeutic benefit.^12,29^ However, ASOs lack cell type specificity and, by targeting a poison exon, may bypass endogenous regulatory mechanisms, leaving open the possibility of ectopic expression of Na_V_1.1. Importantly, this approach requires repeated intrathecal dosing, presenting a significant burden for this patient population.

AAV-delivered zinc finger (ZF)-based modulation represents a one-time treatment approach where the end goal is also upregulation of endogenous *SCN1A* expression.^13^ However, protein-DNA recognition by ZFs is inherently context dependent, which can lead to reduced fidelity and difficult-to-predict off-target activity compared to CRISPR-based systems.^52,53^

A third strategy currently being developed involves split-intein delivery of *SCN1A* cDNA across two separate AAV vectors.^28^ While this approach circumvents the viral genome packaging constraint that precludes delivery of full length *SCN1A* in a single AAV, it increases the risk of inducing non-physiological *SCN1A* expression levels because, unlike gene modulation approaches, it does not interact with the endogenous gene regulatory machinery. Additionally, there are significant regulatory, biodistribution, and manufacturing challenges associated with clinical development of a two-vector system.

In addition to supporting the development of this technology in Dravet syndrome, these findings highlight the broader potential of cell type-selective CRISPR-based transcriptional modulation across genetic diseases. The modular basis of the system comprising the capsid, promoter, effector domain, and gRNA makes it highly adaptable to other tissues, cell types, and target genes. A natural extension of this strategy is to other CNS haploinsufficiencies. For example, *SCN2A*-associated haploinsufficiency has recently been shown to be amenable to CRISPRa-mediated upregulation, with improvement in key phenotypes observed in both human cellular and mouse models.^31^ More broadly, recent advances in capsid engineering, such as the development of variants with enhanced tropism for cardiac and skeletal muscle, will enable application of this approach to haploinsufficiencies affecting peripheral tissues.^54–57^ Finally, this platform could be extended to disorders caused by overexpression or gain-of-function mutations by replacing the transcriptional activator with a repressor domain, such as KRAB or SID, enabling both upregulation and downregulation of gene expression from the same basic framework.^58^

Together, these findings support the continued development of AAV9-E2-dCas9-VP64 (RT101) for Dravet syndrome, and RT101 is currently in preclinical development and is being advanced toward clinical evaluation. More broadly, this work demonstrates that cell type-selective epigenetic modulation has the potential to overcome key safety and efficacy limitations of other genetic medicine technologies, and represents a promising therapeutic strategy for disorders of the CNS and beyond.

## Methods

### gRNA nomination

gRNAs targeting both the mouse and human *SCN1A* genomic loci were nominated using the combined output of multiple gRNA design tools.^59,60^ For mouse-specific gRNAs, nomination was performed on three target regions: from the TSS to 1000 bp upstream for both the distal (“PA”) and proximal (“PB”) *Scn1a* promoters and approximately 200 bp upstream of the E2 enhancer in the first intron (“In”), selected due to proximity of the enhancer and high cross-species conservation. The top 12 mouse gRNAs (based on combined rank) for each promoter region and the top 6 gRNAs for the intronic regions were selected for in vitro screening. For human gRNAs, nomination was performed from 500 bp downstream to 1000 bp upstream of the TSS of the proximal promoter. gRNAs were removed if they had poly-T sequences, high off-target potential, or frequent SNPs. The remaining gRNAs were selected for in vitro screening.

### Recombinant AAV production

rAAV was produced at Franklin Biolabs using standard production methods with triple transfection of the transfer plasmid, AAV9 or PHP.eB rep/cap plasmid, and pHelper in HEK-293T adherent cells. rAAV particles were purified by iodixanol gradients. Titration was performed by ddPCR targeting the dCas9 transgene. For the NHP study, ProHance contrast agent was added to the AAV9 test article prior to injections at a final concentration of 2 mM.

### Luciferase assay

HEK-293T cells (ATCC) were cultured using standard techniques in DMEM+10% FBS. 24 hours after plating in a 96 well plate, cells were transfected with a firefly luciferase reporter plasmid containing the human or mouse *SCN1A* promoter, Renilla luciferase plasmid (internal control), dCas9 plasmid, and gRNA-containing plasmids using X-tremeGene (Sigma) in quadriplicate. 48 hours following transfection, the luciferase assay was carried out according to the manufacturer’s instructions (Dual-Glo Luciferase Assay System, Promega) and luminescence was assessed using a plate reader. Firefly luciferase signal was normalized to Renilla signal and then to the average of non-targeting gRNA controls.

### In vitro transfection and transduction

For transfection experiments to test gRNA activity, Neuro2a (mouse, ATCC), Kelly (human, Sigma), and SK-N-BE(2) (human, ATCC) cells were used. Neuro2a cells were cultured in DMEM (Gibco) +10% FBS and transfected using lipofectamine LTX (Thermo) according to the manufacturer’s instructions. Kelly cells were cultured in RPMI 1640 (Gibco) + 2mM Glutamine + 10% FBS and transfected using lipofectamine LTX (Thermo).

SK-N-BE(2) cells were cultured in DMEM/F12 (Gibco) + 10% FBS and transfected using GeneX Plus (ATCC). For the initial mouse gRNA screen, Neuro2a cells were transfected with a single plasmid containing both the dCas9 transgene and the gRNA sequence. Neuro2a cells were transfected 24 hours after plating and harvested 48 hours following transfection. For the human gRNA screen and testing of the human gRNA in the mouse genome, Kelly, SK-N-BE(2), and Neuro2a cells were transfected 24 hours after plating using a dual vector system in which the gRNA-containing plasmid also contained puromycin resistance. 24 hours post-transfection, puromycin was added to the media. A further 48 hours later, cells were harvested for RNA extraction.

For in vitro AAV transduction experiments, iCell GABANeurons *SCN1A* HZ KO (Fujifilm CDI) were plated in 6 well plates coated with Poly-L-Ornithine (0.01%) and laminin (5ug/mL) and maintained in iCell GABANeuron Maintenance Media + iCell GABANeuron Supplement (Fujifilm CDI). 24 hours after plating, cells were transduced with AAV at the indicated MOIs then cultured for a further 14 days before harvest. Harvested cell pellets were split for RNA and DNA extraction.

### Animals

#### Mouse

Mouse studies were conducted at PsychoGenics, Inc., an AAALAC accredited facility. Procedures were approved by the Institutional Animal Care and Use Committee in accordance with the National Institutes of Health Guide for the Care and Use of Laboratory Animals.

Heterozygous 129S-*Scn1a*^*tm1Kea*^/Mmjax males (The Jackson Laboratory, #024761) were paired with C57BL/6J wild type (WT) female mice procured from The Jackson Laboratory (#000664). At P0, tail samples for all mice were genotyped. Mice were maintained on 12/12 light/dark cycles with temperature between 20 and 23°C and humidity around 50%. Food and water were provided ad libitum. Male and female animals were enrolled in each study. Animals were weighed weekly and survival and health checks were performed twice per day.

#### NHP

NHP studies were conducted at NorthernBio (NB), an AAALAC accredited facility, and complied with all applicable sections of the current version of the Final Rules of the Animal Welfare Regulations (9 CFR), and the Guide for the Care and Use of Laboratory Animals, Institute of Laboratory Animal Resources, Commission on Life

Sciences, National Research Council.

AAV9-seronegative cynomolgus macaques aged 2-3 years old originating from Mauritius were socially housed with environmental enrichment, separated by sex and group. Animals were fed a facility approved diet (Teklad Diets, Madison, WI) and water (City of Muskegon, MI) was provided ad libitum by an automatic watering system. Cage side clinical observations were conducted throughout the study, with detailed clinical observations performed on days 1, 3, and 4 following dosing, and once weekly thereafter. Body weights were collected weekly.

### Intracerebroventricular injection

#### Mouse

Cryoanesthetized animals were dosed via bilateral intracerebroventricular injection (ICV) on postnatal day 1, with a dose volume of 3 µl per hemisphere. Treatments were then administered using 10 µl Hamilton Syringes (1701 RN Model, Cat# 7653-01) and Hamilton 30½ g needles (point style 4, bevel 12°, Cat# 7803-07). Following injection, animals were placed on a warm (~36°C) heating pad. Pups were returned to the dam once all animals from the litter were dosed and recovered from cryoanesthesia.

#### NHP

Animals were anesthetized prior to injection following NB standard operating procedures. The scalp was shaved, and the animals were mounted in a stereotaxic frame. Baseline MRI images were used to establish target location. An incision was made and a single hole was drilled through the skull over the target location. A modified ClearPoint cannula (ClearPoint Neuro, Solana Beach, CA) was primed with the AAV9 test article and loaded into the frame. The dura was opened and the cannula was advanced to the appropriate depth using MRI guidance. The preloaded syringe was placed into an infusion pump, and dosing material was delivered at a rate of 0.1 mL/minute for a duration of 10 minutes (1 mL delivered in total). Contrast agent (Prohance, 2 mM) included in the test article formulation was used to confirm correct cannula placement.

### Hyperthermia-induced seizure assessment

Mice were placed in a temperature control module with an anal temperature probe and allowed to acclimate to the chamber for five minutes, followed by a two minute probe stabilization period during which the baseline body temperature for each mouse was recorded. The temperature was then increased by approximately 0.5°C every two minutes using a heat lamp until the first visible tonic-clonic seizure occurred, at which point the animal was transferred to a cooling chamber. Mice were considered not to have seized if they reached 43ºC without a tonic-clonic seizure. Body temperature was recorded continuously, starting at probe insertion and concluding at probe removal. Animals subjected to the hyperthermia protocol were sacrificed for tissue collection following the conclusion of the assay.

### Tissue collection

#### Mouse

Samples were collected for molecular analysis by anesthetizing animals with isoflurane followed by transcardial perfusion with artificial cerebrospinal fluid (ACSF) to remove blood cells. Whole brains were removed, weighed, coronally sectioned into 1 mm sections using a brain block, and subsequently snap-frozen in well plates on dry ice. All other tissue samples were snap-frozen in microcentrifuge tubes in liquid nitrogen.

For histological analysis, mice were anesthetized with isoflurane and transcardially perfused with PBS followed by 4% paraformaldehyde (PFA). Brains were post-fixed in 4% PFA overnight and transferred to PBS.

#### NHP

Animals were sacrificed for tissue collection in accordance with NB euthanasia and necropsy protocols. Whole brains were collected and sectioned into 4 mm thick coronal sections.Interleaving slabs were processed for histological assessment by post-fixation in 4% PFA followed by transfer to PBS after 24 hours, or 4 mm tissues punches were taken from pre-defined brain regions and flash frozen for bioanalysis. The spinal cord, dorsal root ganglion and peripheral organs were removed, with separate samples either post-fixed in 4% PFA followed by transfer to PBS for histology, or flash-frozen for bioanalysis.

### Nucleic acid extraction

#### In vitro samples

For all in vitro experiments, RNA extraction was performed using the mirVana MagMAX Total RNA Isolation Kit including Turbo DNAse treatment per kit instructions and, where applicable, MagMAX DNA Multi-Sample Ultra 2.0 Kit.

#### Mouse

For mouse brain samples, four 2 mm diameter biopsy punches were taken from the somatosensory cortex from 1 mm thick brain slices and homogenized using a motorized pestle. For all other mouse tissues, 3 mm zirconium beads were used to homogenize tissue with a Mini-Beadbeater (BioSpec).

#### NHP

Flash-frozen brain tissue punches were homogenized with 3 mm zirconium beads in a Mini-Beadbeater (BioSpec). For all other tissues, approximately 20 mg of frozen tissue was homogenized with 3 mm zirconium beads in a Mini-Beadbeater.

DNA and RNA were extracted from homogenized mouse and NHP tissue using the AllPrep DNA/RNA Mini kit (Qiagen) according to the manufacturer’s instructions. RNA was immediately DNase-treated with Turbo DNase (Invitrogen) for 30 minutes following manufacturer’s instructions. DNA and RNA concentrations were quantified via NanoDrop.

### Vector biodistribution analysis

Genomic DNA concentrations were normalized across samples and then vector genome and reference genome abundance was quantified by digital droplet polymerase chain reaction (ddPCR). Droplets were generated with an AutoDG droplet generator (Bio-Rad), amplification was performed in a C1000 Touch Thermal Cycler (Bio-Rad), and droplets were read with a QX600 droplet reader (Bio-Rad). For mouse samples, Ptbp2 was used to quantify reference genomes.^61^ For NHP samples, HBE1 was used to quantify reference genomes.^62^ For NHP samples, 250 ng of genomic DNA was digested by SmaI for 1 hour prior to ddPCR to fragment vector genome concatemers.^63^ To quantify vector genomes, custom assays were designed that target the regulatory regions present in our vector. For Figure 1, a probe-based assay that targets the gRNA promoter was used. For Figure 4, a probe-based assay that spans the gRNA promoter and gRNA scaffold was used. For Figure 5, a probe-based assay that targets our dCas9 promoter was used.

### RNA expression analysis

Reverse transcription of total RNA was performed using Ultra SuperMix (QuantaBio) following manufacturer’s instructions on 100 ng (in vitro samples), 500 ng (mouse), or 200 ng (NHP) of RNA. For Neuro2a, Kelly, and SK-N-BE(2) experiments, qPCR was performed on a QuantStudio 6. Neuro2a qPCR was performed using SYBR-based assays, where expression of *Scn1a* (FWD: CTCTTCCCCGCATCAGTCTC, REV: TTCGGGGAACGAACAGTGAG) was normalized to expression of *Actb*.^*62*^ Kelly and SK-N-BE(2) qPCR was performed using probe-based assays, where *SCN1A* expression (Hs.PT.58.15238557, IDT) was normalized to the geometric mean of the expression of *RPS18* (Hs.PT.58.14390640, Bio-Rad), *HPRT1* (qHsaCIP0030549, Bio-Rad), and *TBP* (qHsaCIP0036255, Bio-Rad).

RNA expression analysis from iCell GABA, mouse, and NHP samples were performed via ddPCR as described for vector biodistribution analysis. To quantify dCas9 and gRNA expression, custom probe-based assays were designed. A probe-based assay for dCas9-VP64 was used for Figure 1, and a probe-based assay for the 3’ end of dCas9 was used for Figures 3 through 5. For iCell GABA ddPCR, expression of dCas9, gRNA, *SCN1A (*Hs.PT.58.15238557, IDT), *SCN7A* (Hs.PT.58.27029586, IDT), *GALNT3* (Hs.PT.58.19802430, IDT), *SCN9A* (Hs.PT.58.20989243, IDT), *CSRNP3* (Hs.PT.58.27479232, IDT), *TTC21B* (Hs.PT.58.4061727, IDT), and *SCN2A* (Hs.PT.58.3174775, IDT) was normalized to the geometric mean of the expression of *RPS18* (Hs.PT.58.14390640, IDT) and *TBP* (dHsaCPE55203391, Bio-Rad). For mouse samples, expression of dCas9 and the WT *Scn1a* allele (Mm.PT.58.13867113, IDT) was normalized to expression of *Hprt* (Mm.PT.58.32092191, IDT).

### Genomic off-target analysis

RNA samples were submitted to Azenta Life Sciences for library preparation using strand-specific RNA-seq with poly(A) selection and sequencing (2 × 150 bp; ~30 million read pairs per sample). Reads were aligned to the human reference genome (hg38) using STAR (v2.5.2b), and duplicate reads were removed using Samtools (v1.21). Gene-level counts were generated with the Rsubread package (v2.16.1) using the Gencode v47 basic annotation set. Differential expression analysis was performed with DESeq2 (v1.44.0) for each cell line, comparing *SCN1A*-g10 to a non-targeting control (NC01). Genes with normalized counts ≥10 in at least two samples were retained for analysis. Off-target sites were predicted using Cas-OFFinder^42^ allowing up to 5 mismatches or 1 DNA/RNA bulge plus 1 mismatch. Genes with transcription start sites within 500 kb of predicted off-target sites (based on the closest annotated TSS for each gene in the Gencode v47 basic annotation set) were identified using bedTools (v2.31.1).^64^ Among these, significantly upregulated genes (log2FC > 0.3, adjusted P ≤ 0.05; Wald test) were considered potential off-targets.

### Na_**V**_1.1 protein Mesoscale Diagnostics electrochemiluminescence assay

Frozen coronal slices of mouse brain that include somatosensory cortex were transferred to IP buffer (Thermo Fisher) with protease and phosphatase inhibitors (Sigma), and then homogenized with 3 mm zirconium beads in a Mini-Beadbeater (BioSpec). Total protein concentrations were measured with the Pierce Dilution-Free BCA Rapid kit (Thermo Fisher), and diluted to 3 mg/mL.

To prepare for its use as a capture antibody in our Mesoscale Diagnostics (MSD) assay, a mouse Anti-Na_V_1.1 antibody (Antibodies Inc) was biotinylated in house with the EZ-Link Sulfo-NHS-LC-Biotinylation Kit (Thermo) following manufacturer’s instructions. A standard curve was prepared using serial dilutions of wild-type brain lysate prepared from littermate controls. Briefly, small spot streptavidin plates (MSD) were blocked with 1% Blocker B (MSD) in TBS-T for 1 hour. Plates were washed and incubated with the biotinylated antibody for 1 hour. After washing, samples were added at 3 mg/mL and incubated overnight at 4°C. The following morning, the plates were washed and incubated with a rabbit anti-Na_V_1.1 detection antibody (Alomone, Inc) for one hour. After an additional wash step, MSD SULFO-tag antibody was applied for 1 hour prior to reading in MSD Gold Read Buffer with a MESO QuickPlex SQ 120MM instrument. Electrochemiluminescence (ECL) values were interpolated against the WT standard curve, and Na_V_1.1 protein in each sample is reported as a percentage of WT protein.

### Immunohistochemistry

#### Mouse

Fixed brains were bisected along the longitudinal fissure, embedded in 4% agarose, and sectioned sagittally at 50 µm on a Leica VT1000 S vibratome. Free-floating sections were quenched with 3% H_2_O_2_, blocked in 10% normal goat serum (NGS) in PBS.

#### NHP

Fixed 4 mm brain slices were trimmed to appropriate size and washed with a graded ethanol and xylene series followed by paraffin embedding. Formalin-fixed paraffin-embedded tissue was sectioned at 5 µm and mounted onto SuperFrost Plus slides. Slides were baked at 60°C for 1 hour, cooled, and deparaffinized through xylene and a graded ethanol series (100%, 95%, 70%) before rehydration. Endogenous peroxidase was quenched with H_2_O_2_ (Tyramide SuperBoost Kit, Thermo Fisher, B40944), antigen retrieval was performed in citrate buffer (IHCworld, IW-1100) at 99–100°C for 15 minutes, and sections were blocked in 10% NGS.

Sections were incubated overnight at 4°C with primary antibodies (rabbit monoclonal anti-SpCas9, Abcam, ab203943, 1:100; guinea pig anti-parvalbumin, Swant, GP72, 1:2000 for mouse, 1:1000 for NHP) in 10% NGS. Sections were then incubated with HRP-conjugated goat anti-rabbit secondary (Tyramide SuperBoost Kit, Thermo Fisher, B40944) followed by tyramide signal amplification with Alexa Fluor 594 tyramide reagent. Sections were then incubated with donkey anti-guinea pig Alexa Fluor 488 (Jackson ImmunoResearch, 706-545-148; 1:500) in 0.3% Triton X-100/5% NDS/PBS, mounted onto slides, and coverslipped with Fluoromount-G with DAPI.

Fluorescence imaging from mouse tissue was acquired using an FV 4000 confocal microscope (FLUOVIEW), while imaging of NHP tissue was performed using an AxioObserver 7 inverted microscope (Zeiss). Intensity and exposure settings were held constant across all animals. Tiled images were quantified using Photoshop (Adobe). Brightness and contrast were adjusted per channel and held constant across all images. Cells were counted manually using the Photoshop count tool. A cell was considered positive if the signal was above background fluorescence. Sensitivity for PV-positive cells was calculated as the ratio of co-labeled cells to the total number of PV-positive cells. Specificity for PV-positive cells was calculated as the ratio of co-labeled cells to the total number of dCas9-positive cells.

### In Situ Hybridization for Vector DNA Detection

In situ hybridization was performed using the RNAscope Multiplex Fluorescent Reagent Kit v2 (ACD Bio, 323100) adapted for DNA detection. Sections were mounted onto SuperFrost Plus slides, baked at 60°C, post-fixed in 4% PFA, quenched with hydrogen peroxide, and subjected to target retrieval by briefly immersing slides in 100ºC water followed by transfer to target retrieval reagent (provided by kit). An oligonucleotide probe targeting the non-transcribed DNA sequence of the Sa-dCas9-VP64 cassette (ACD Bio, 1163241-C1) was hybridized at 40°C for 2 hours. After overnight storage in 2X SSC buffer (cat#: AAJ60839-K2), signal was amplified sequentially (AMP1–3) and detected with Opal 570 fluorophore (Akoya Biosciences, FP1488001KT; 1:1500) via HRP/TSA. Slides were coverslipped with Fluoromount-G with DAPI.

60x tiled images were acquired of somatosensory sections using an FV 4000 confocal microscope (FLUOVIEW). Acquisition settings were determined at the time of imaging, and all imaging and post-image processing parameters were held constant across animals and groups to enable direct comparison.

## Acknowledgements

The authors thank all current and former employees of Regel Therapeutics for their contributions and helpful discussions which made this work possible including: Stephen Farr, Ryan Ziffra, Jeanne Duong, Mia Brakebill, Jeffrey Rubin, Raul Morales, Vaishali Muralidaran, Molly Gordon, Tasneem Rinvee, Craig Ennis, Megan Rowton, Nick Aguirre, Chris Streilein, David Saul, Bersh Marius, Andrea Gorodezky, Benjamin Cook, and Benjamin Whiting.

We also thank the teams at PsychoGenics, NorthernBio, and Franklin Biolabs for conducting the mouse and non-human primate in vivo studies and AAV manufacturing, respectively. *Scn1a*^tm1Kea^ mice were provided by Vanderbilt University. This work was funded by Regel Therapeutics, Inc.

## Author Contributions

P.W.E.S., N.F.T., and K.C.A. conceived and designed the studies and wrote the original manuscript draft. P.W.E.S. and K.C.A. coordinated external studies conducted at contract research organizations. P.W.E.S., N.F.T., R.M.G., O.S., T.N., J.R., L.C., A.R.F., J.E., M.F., and Y.H. conducted experiments and analyzed data. C.H. coordinated AAV production and contributed to study design discussions and data interpretation. J.D. N.M., and O.D. conceived the original therapeutic strategy. J.D. and K.C.A. supervised the work. J.D., R.M.G., O.S., T.N., J.R., and A.R.F. reviewed and edited the manuscript. All authors reviewed and approved the final manuscript.

## Supplementary Information

**Supplementary Fig. S1.**
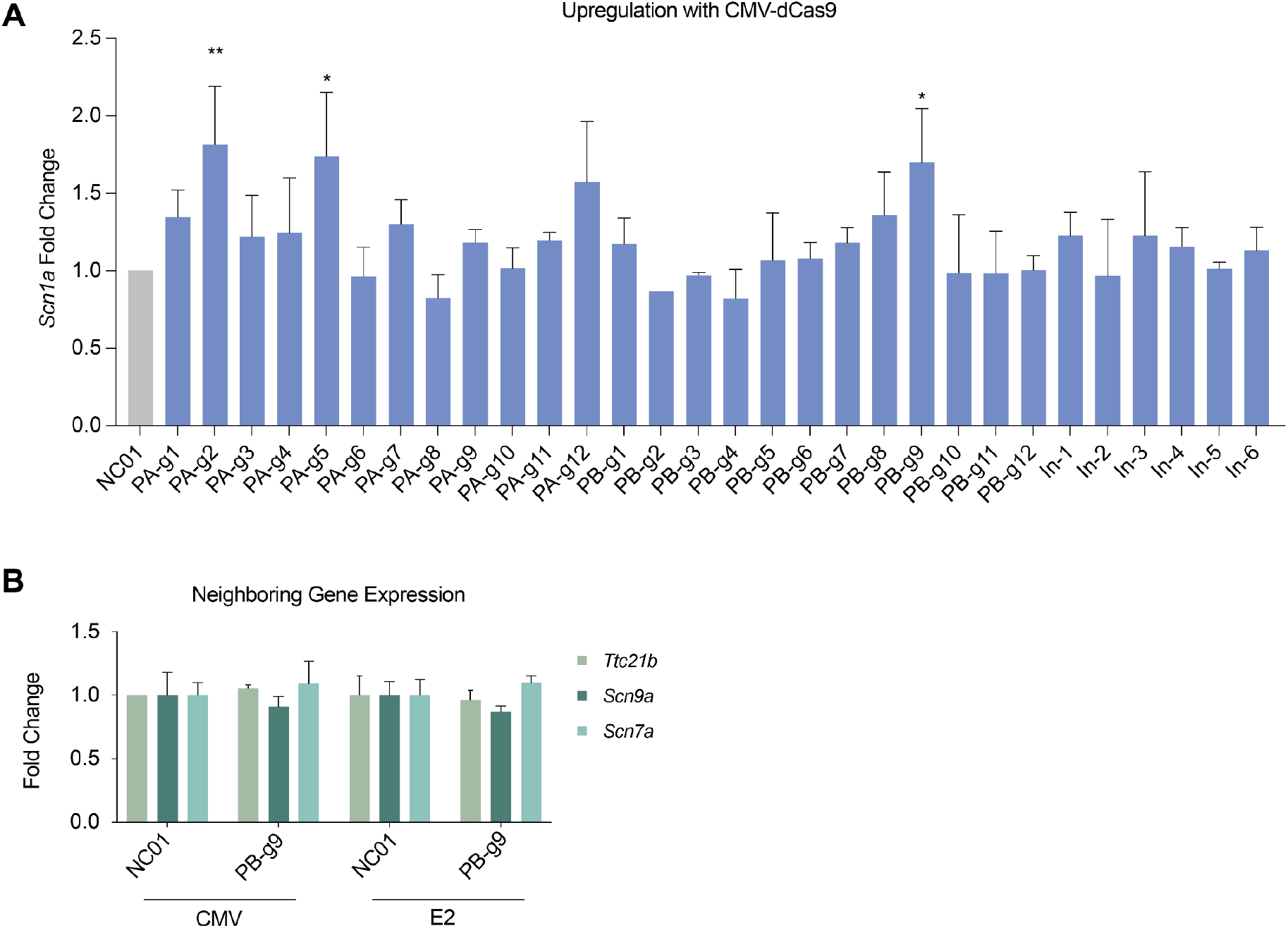
Identification of a gRNA targeting the mouse *Scn1a* locus. **(A)** Upregulation of *Scn1a* in vitro in Neuro2a cells measured by qPCR following transfection with a CMV-dCas9-hU6-gRNA all-in-one plasmid. Expression normalized to *Actb* and shown as fold change relative to non-targeting gRNA (NC01). n=2-4 per gRNA. Statistical significance determined by one-way ANOVA with multiple comparisons relative to NC01, PA-g2: p=0.0031; PA-g5: p=0.0101; PB-g9: p=0.0186. **(B) E**xpression of genes neighboring *Scn1a* relative to NC01 following transfection with either CMV-dCas9-hU6-*Scn1a*-PB-g9 (CMV) or E2-dCas9-hU6-*Scn1a*-PB-pg9 (E2). n=2. No statistically significant differences relative to NC01 were found (one-way ANOVA with Šídák’s multiple comparisons).

**Supplementary Fig. S2.**
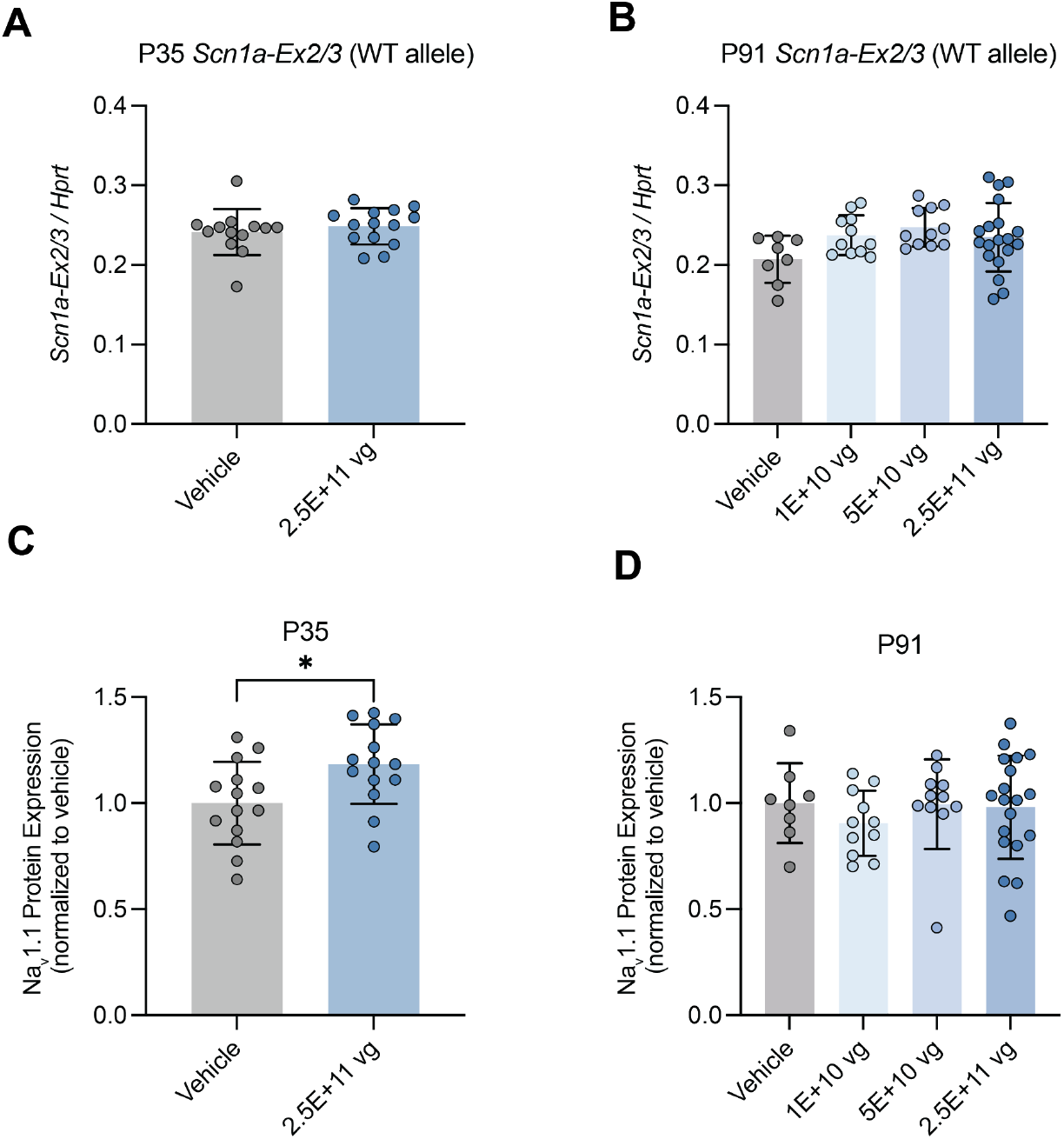
*Scn1a* mRNA and Na_V_1.1 protein expression following AAV9-E2-dCas9-VP64 injection in *Scn1a*^+/-^ mice. **(A)** *Scn1a* mRNA is not significantly upregulated by *Scn1a* PB-g9 at P35. *Scn1a* cp/uL are normalized to Hprt cp/uL. Each point represents one mouse somatosensory cortex. Each bar represents the mean of all mice treated with the indicated dose, error bars represent standard deviation. Vehicle: n=13; 2.5E+11: n=14. **(B)** *Scn1a* mRNA is not significantly upregulated by *Scn1a* PB-g9 at P91. *Scn1a* cp/uL are normalized to Hprt cp/uL. Each point represents one mouse somatosensory cortex. Each bar represents the mean of all mice treated with the indicated dose, error bars represent standard deviation. Vehicle: n=8; 1.0E+10: n=11; 5.0E+10: n=11; 2.5E+11: n=19. **(C)** Na_V_1.1 is upregulated by AAV9-E2-dCas9-VP64 with *Scn1a* PB-g9 in *Scn1a*^+/-^ mice at P35. Na_V_1.1 levels are shown as a percentage of Na_V_1.1 measured in littermate WT controls. Each point represents one mouse somatosensory cortex. Each bar represents the mean of all mice treated with the indicated dose, error bars represent standard deviation. *p=0.0174, Holm-Šídák’s multiple comparisons test following one-way ANOVA. Vehicle: n=14; 2.5E+11: n=14. **(D)** Na_V_1.1 is not significantly upregulated by AAV9-E2-dCas9-VP64 with *Scn1a* PB-g9 in *Scn1a*^+/-^ mice at P91. Na_V_1.1 levels are shown as a percentage of Na_V_1.1 measured in littermate WT controls. Each point represents one mouse somatosensory cortex. Each bar represents the mean of all mice treated with the indicated dose, error bars represent standard deviation. Vehicle: n=8; 1.0E+10: n=11; 5.0E+10: n=11; 2.5E+11: n=19.

**Supplementary Fig. S3.**
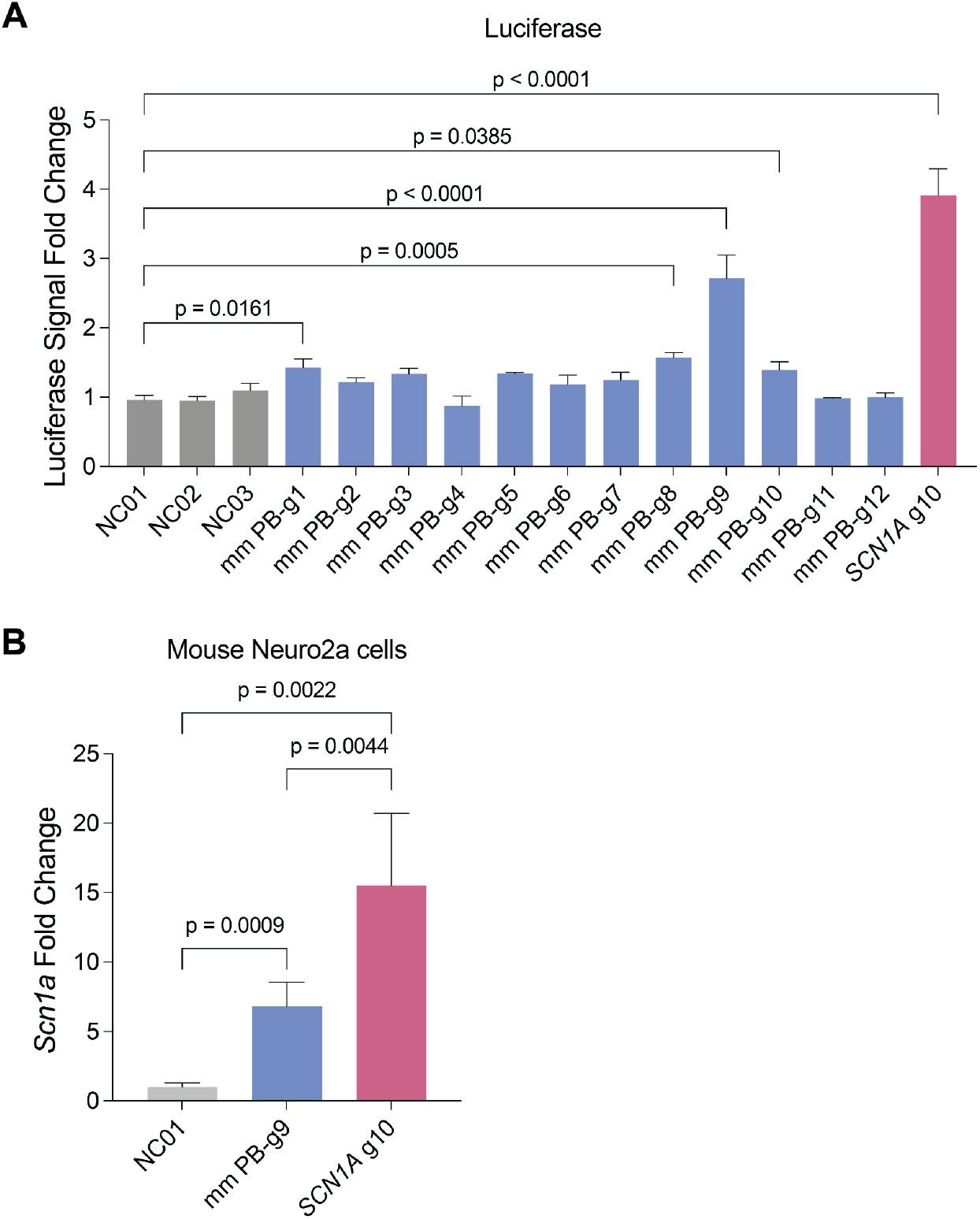
Assessment of *SCN1A*-g10 gRNA activity in the mouse genome in vitro. **(A)** Luciferase assay (HEK-293T cells) measuring upregulation of luciferase under the control of the mouse *Scn1a* promoter. Mouse-specific gRNAs (mmPX-X) were compared to the *SCN1A*-g10 gRNA identified in human cells with sequence conservation in mice. Normalized luminescence levels shown as fold change relative to the average of non-targeting gRNA controls (NC0X). n=3 per gRNA. Statistical significance determined by one-way ANOVA with Šídák’s multiple comparisons relative to NC01, p-values indicated on figure. **(B)** Assessment of endogenous *Scn1a* upregulation in Neuro2a cells with the original, mouse-specific gRNA tested in vivo (mmPB-g9, Figure 1) and the gRNA identified with sequence conservation in both the human and mouse genomes (*SCN1A*-g10). Neuro2a cells were transfected with the two-vector system described in Figure 2 and upregulation was measured by qPCR. Statistical significance determined by one-way ANOVA with Tukey’s multiple comparisons, p-values indicated on figure.

